# Polynomial-Time Statistical Estimation of Species Trees under Gene Duplication and Loss

**DOI:** 10.1101/821439

**Authors:** Brandon Legried, Erin K. Molloy, Tandy Warnow, Sébastien Roch

**Author notes:** {, }.

## Abstract

Phylogenomics—the estimation of species trees from multilocus datasets—is a common step in many biological studies. However, this estimation is challenged by the fact that genes can evolve under processes, including incomplete lineage sorting (ILS) and gene duplication and loss (GDL), that make their trees different from the species tree. In this paper, we address the challenge of estimating the species tree under GDL. We show that species trees are *identifiable* under a standard stochastic model for GDL, and that the polynomial-time algorithm ASTRAL-multi, a recent development in the ASTRAL suite of methods, is *statistically consistent* under this GDL model. We also provide a simulation study evaluating ASTRAL-multi for species tree estimation under GDL. All scripts and datasets used in this study are available on the Illinois Data Bank: https://doi.org/10.13012/B2IDB-2626814_V1.

## 1 Introduction

Phylogeny estimation is a statistically and computationally complex estimation problem, due to heterogeneity across the genome resulting from processes such as incomplete lineage sorting (ILS), gene duplication and loss (GDL), rearrangements, gene flow, horizontal gene transfer, introgression, etc. [23]. Much is known about the problem of estimating species trees in the presence of ILS, as modelled by the Multi-Species Coalescent (MSC) [19,37]. For example, because the most probable unrooted tree for every four species is the species tree on those species [1], the unrooted species tree topology is identifiable under the MSC from its gene tree distribution, and quartet-based species tree estimation methods that operate by combining gene trees (such as BUCKy-pop [20] and ASTRAL [25,27,44]) are statistically consistent estimators of the unrooted species tree topology (i.e., as the number of sampled genes increases, almost surely the tree returned by these methods will be the true species tree). It is also known that concatenation (whether partitioned or unpartitioned) is not statistically consistent, and can even be positively misleading (i.e., converge to the wrong tree as the number of loci increases) [34,32]. In general, establishing whether a method is statistically consistent or not is important for understanding its performance guarantees.

Yet, correspondingly little has been established about species tree estimation in the presence of GDL. For example, although likelihood-based approaches for species tree estimation have been developed (e.g., PHYLDOG [7]), they have not been established to be statistically consistent. Key to understanding the performance of species tree estimation under GDL is whether the species tree topology itself is identifiable from the distribution it defines on the gene trees it generates. However, since gene trees can have multiple copies of each species when gene duplication occurs, this question can be formulated as: “Is the species tree identifiable from the distribution on MUL-trees?”, where a MUL-tree is a tree with potentially multiple copies of each species.

In this paper, we prove that unrooted species tree topologies are identifiable from the distribution implied on MUL-trees (Section 3) under the simple GDL model of [2]. Furthermore, we prove that the polynomial-time method ASTRAL-multi [29], a recent variant of ASTRAL designed to enable analyses of datasets with multiple individuals per species, is statistically consistent under this model (Section 3). We then present an experimental study evaluating ASTRAL-multi on 16-taxon datasets simulated under the DLCoal model (a unified model of GDL and ILS) [30]; the results of this study show that when given a sufficiently large number of genes, ASTRAL-multi is competitive with other methods (e.g., Dup-Tree [4], MulRF [9], and ASTRID-multi [39], the implementation of ASTRID for multi-allele datasets) that also estimate species trees from MUL-trees (Section 4). We conclude with remarks about future work and implications for large-scale species tree estimation (Section 5).

## 2 Species tree estimation from gene families

Our input is a collection 𝒯 of gene trees representing the inferred evolutionary histories of gene families. In the presence of gene duplication and loss events, such gene trees may be multi-labeled trees (MUL-trees), meaning that the same species label may be assigned to several gene copies. Our goal is to reconstruct a species tree *T* over the corresponding set *S* of species.

### ASTRAL

We provide theoretical guarantees and empirically validate an approach based on ASTRAL [25] in its variant for multiple alleles [29], which we refer to as ASTRAL-multi. Following [12], the input consists of unrooted MUL-trees 𝒯 from all gene families, where copies of a gene in a species are treated as multiple alleles within the species.

ASTRAL-multi proceeds as follows. Let *S* be the set of *n* species and let *R* be the set of *m* individuals. The input are the gene trees 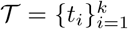, where *t*_*i*_ is labeled by individuals *R*_*i*_ ⊆ *R*. For any (unrooted) species tree 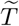 labeled by *S*, an extended species tree 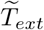 labeled by *R* is built by adding to each leaf of 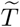 all individuals corresponding to that species as a polytomy. The quartet score of 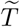 with respect to 𝒯 is then

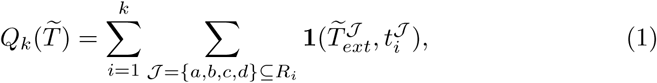

where **1**(*T*_1_, *T*_2_) is the indicator that *T*_1_ and *T*_2_ agree and 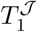 is the restriction of *T*_1_ to individuals 𝒥. Run in its *exact* version (i.e., an unrooted species tree that maximizes the quartet score), ASTRAL-multi is guaranteed to find an optimal solution, but can use exponential time. The *default* mode, which runs in polynomial time, uses dynamic programming to solve a constrained version of the problem, requiring that the output tree draw its bipartitions from a set *Σ* of bipartitions that ASTRAL computes on the input, where *Σ* by construction includes all the bipartitions on *S* that occur in any gene tree in 𝒯.

## 3 Theoretical results

In this section, we provide theoretical guarantees for the reconstruction algorithm discussed in Section 2. Specifically, we establish statistical consistency under a standard model of GDL [2]. First we show that the species tree is identifiable.

### 3.1 Gene duplication and loss model

We assume in this section that gene tree heterogeneity is due exclusively to GDL (and so no ILS) and that the true gene trees are known. That is, there is no gene tree estimation error (GTEE).

#### Birth-death process of gene duplication and loss

The rooted *n*-species tree *T* = (*V, E*) has vertices *V* and directed edges *E* with lengths (in time units) *η* that depend on the edge. For ease of presentation, we assume that there is a single copy of each gene at the root of *T* and that the rates of duplication *λ* and loss *µ* are fixed throughout *T* (although our proofs do not use these assumptions). Each gene tree is generated by a top-down birth-death process within the species tree. That is, on each edge, each gene copy independently duplicates at exponential rate *λ* and is lost at exponential rate *µ*; at speciation events, each gene copy bifurcates and proceeds similarly in the descendant edges. Each duplication is indicated in the gene tree by a bifurcation. The resulting gene tree is then pruned of lost copies to give the observed unrooted gene tree *t*_*i*_. The gene trees 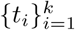 are assumed independent and identically distributed. See more details in [2].

### 3.2 Identifiability of the species tree under the GDL model

We first show that the unrooted species tree is identifiable from the distribution of MUL-trees 𝒯 under the GDL model over *T*. That is, that two distinct unrooted species trees necessarily produce different gene tree distributions.

We begin with a quick proof sketch. The idea is to show that, for each 4-tuple of species 𝒬 = {*A, B, C, D*}, the corresponding species quartet topology can be identified by taking an independent uniform random gene copy in each species in 𝒬 and showing that the quartet topology consistent with the species tree is most likely to result in the gene tree restricted to these copies. It should be noted that the proof is not as straightforward as it is under the multispecies coalescent [1], as we explain next. Assume the species tree restricted to 𝒬 is ((*A, B*), (*C, D*)), let *R* be the most recent common ancestor of 𝒬 in *T*, and let *a, b, c, d* be random gene copies in *A, B, C, D* respectively.

– When all ancestral copies of *a, b, c, d* in *R* are distinct, by symmetry *all quartet topologies are equally likely*. The *ancestral copy of x in R* is the vertex of the gene tree that is ancestral to *x* and corresponds to a speciation event at node *R* of the species tree.
– When the ancestors of *a* and *b* (or *c* and *d*) in *R* are the same, the *species quartet topology results*.
– *However*, there are further cases. For example, if the ancestors of *a* and *c* in *R* coincide while being distinct from those of *b* and *d*, then the resulting quartet topology *differs* from that of the species tree.

Hence, one must carefully account for all possible cases to establish that the species quartet topology is indeed likeliest, which we do next. Our argument relies primarily on the symmetries (i.e., exchangeability) of the process.

#### Theorem 1 (Identifiability).

*Let T be a species tree with n* ≥ 4 *leaves. Then T, without its root, is identifiable from the distribution of MUL-trees 𝒯 under the GDL model over T*.

*Proof*. It is known that the unrooted topology of a species tree is defined by its set of quartet trees [3]. Let 𝒬 = {*A, B, C, D*} be four distinct species in *T* and let *T*^𝒬^ be the species tree restricted to 𝒬. Assume without loss of generality that the corresponding unrooted quartet topology is *AB*|*CD*. Let *t* be a MUL-tree generated under the GDL model over *T* and let *t*^𝒬^ be its restriction to the gene copies from species in 𝒬. Conditioning on having at least one gene copy in the species 𝒬, independently pick a uniformly random gene copy *a, b, c, d* in species *A, B, C, D* respectively and let *q* be the corresponding quartet topology under *t*^𝒬^. We show that the most likely outcome is *q* = *ab*|*cd*. There are two cases: *T*^𝒬^ is 1) balanced or 2) a caterpillar.

In case 1), let *R* be the most recent common ancestor of 𝒬 in *T* and let *I* be the number of gene copies exiting (forward in time) *R*. By the law of total probability, **P**′[*q* = *ab*|*cd*] = **E**′[**P**′_*I*_[*q* = *ab*|*cd*]], where the primes indicate that we are conditioning on having at least one gene copy in each species in 𝒬 and the subscript *I* indicates conditioning on *I*. So it suffices to prove

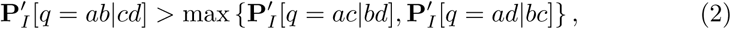

almost surely. Let *i*_*x*_ ∈ {1,…, *I*} be the ancestral lineage of *x* ∈ {*a, b, c, d*} in *R*. Then

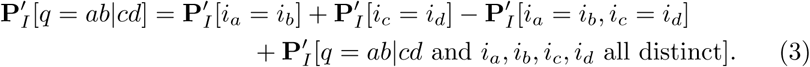

On the other hand,

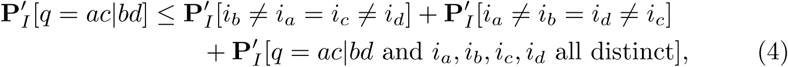

and similarly for **P**′_*I*_[*q* = *ac*|*bd*], where note that we double-counted the case *i*_*a*_ = *i*_*c*_ ≠ *i*_*d*_ = *i*_*b*_ to simplify the expression. By symmetry of the GDL process above *R* (which holds under **P**′_*I*_), the last term on the RHS of (3) and (4) are the same. The same holds for the first two terms on the RHS of (4) this time by the independence and exchangeability of the pairs (*i*_*a*_, *i*_*b*_) and (*i*_*c*_, *i*_*d*_) under **P**′_*I*_, which further implies

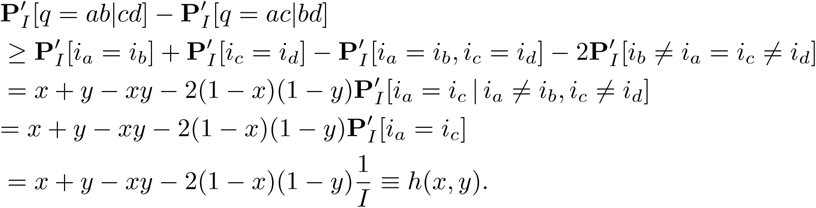

where *x* = **P**′_*I*_ [*i*_*a*_ = *i*_*b*_] and *y* = **P**′_*I*_ [*i*_*c*_ = *i*_*d*_].

For fixed *y, h*(*x, y*) is linear in *x* and *h*(1, *y*) = 1. So *h*(·, *y*) achieves its minimum at the smallest value allowed for *x*. The same holds for *y*. Intuitively, *i*_*a*_ and *i*_*b*_ are “positively correlated” so *x* ≥ 1*/I*. We prove this formally next.

#### Lemma 1.

*Almost surely, x, y* ≥ 1*/I*.

*Proof*. For *j* ∈ {1,…, *I*}, let *N*_*j*_ be the number of gene copies at the most recent common ancestor *R*′ of *A* and *B* that descend from copy *j* in *R*. Upon conditioning on (*N*_*j*_)_*j*_, the choice of *a* and *b* is independent, with *i*_*a*_ and *i*_*b*_ being picked proportionally to the corresponding *N*_*j*_’s (i.e., the gene copies in *R*′ are equally likely to have given rise to *a*). By the law of total probability and the fact that the quadratic mean is greater than the arithmetic mean,

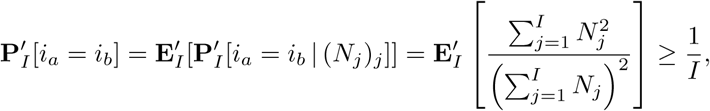

and similarly for **P**′_*I*_ [*i*_*c*_ = *i*_*d*_]. □

Returning to the proof of the theorem, evaluating *h* at *x, y* = 1*/I* gives

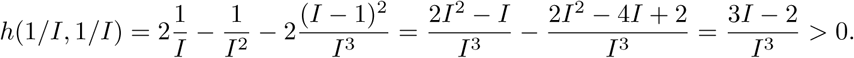

That establishes (2) in case 1), which implies

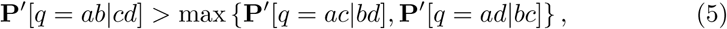

as desired. The proof in case (2) can be found in the appendix. □

As a direct consequence of our identifiability proof, it is straightforward to establish the statistical consistency of the following pipeline, which we refer to as ASTRAL/ONE (see also [12]): for each gene tree *t*_*i*_, pick in each species a random gene copy (if possible) and run ASTRAL on the resulting set of modified gene trees 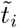. The proof can be found in the appendix.

#### Theorem 2 (Statistical Consistency: ASTRAL/ONE).

*ASTRAL/ONE is statistically consistent under the GDL model. That is, as the number of input gene trees tends toward infinity, the output of ASTRAL/ONE converges to T almost surely, when run in exact mode or in its default constrained version*.

### 3.3 Statistical consistency of ASTRAL-multi under GDL

The following consistency result is not a direct consequence of our identifiability result, although the ideas used are similar.

#### Theorem 3 (Statistical Consistency: ASTRAL-multi).

*ASTRAL-multi, where copies of a gene in a species are treated as multiple alleles within the species, is statistically consistent under the GDL model. That is, as the number of input gene trees tends toward infinity, the output of ASTRAL-multi converges to T almost surely, when run in exact mode or in its default constrained version*.

*Proof*. First, we show that ASTRAL-multi is consistent when run in exact mode. The input are the gene trees 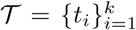 with *t*_*i*_ labelled by individuals (i.e., gene copies) *R*_*i*_ ⊆ *R*. Then the quartet score of 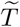 with respect to 𝒯 is given by (1). For any 4-tuple of gene copies 𝒥 = {*a, b, c, d*}, we define *m*(𝒥) to be the corresponding set of species. It was proved in [29] that those 𝒥’s with fewer than 4 species contribute equally to all species tree topologies. As a result, it suffices to work with a modified quartet score

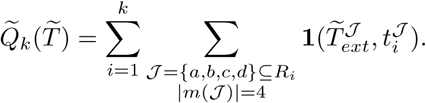

By independence of the gene trees (and non-negativity), 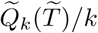 converges almost surely to its expectation simultaneously for all unrooted species tree topologies over *S*.

The expectation can be simplified as

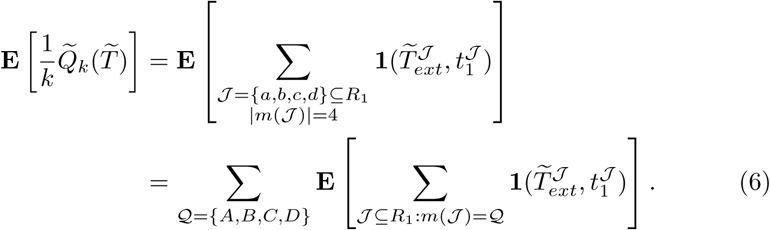

Let 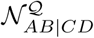 (respectively 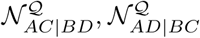) be the number of choices consisting of one gene copy in *t*_1_ from each species in 𝒬 whose corresponding restriction 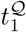 agrees with *AB*|*CD* (respectively *AC*|*BD, AD*|*BC*). Then each summand in (6) may be written as 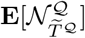. We establish below that this last expression is maximized at the true species tree *T*^𝒬^, that is,

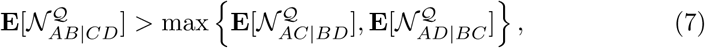

when (without loss of generality) *T*^𝒬^ = *AB*|*CD*. From (6) and the law of large numbers, it will then follow that almost surely the quartet score is eventually maximized by the true species tree as *k* → +∞.

It remains to establish (7). Fix 𝒬 = {*A, B, C, D*} a set of four distinct species in *T*. Assume that the corresponding unrooted quartet topology in *T* is *AB*|*CD*. Let *t*_1_ be a MUL-tree generated under the GDL model over *T*. Again, there are two cases: *T*^𝒬^ is 1) balanced or 2) a caterpillar.

In case 1), let *R* be the most recent common ancestor of 𝒬 in *T* and let *I* be the number of gene copies exiting (forward in time) *R*. For *j* ∈ {1,…, *I*}, let 𝒜_*j*_ be the number of gene copies in *A* descending from *j* in *R*, and similarly define ℬ_*j*_, 𝒞_*j*_ and 𝒟_*j*_. By the law of total probability, 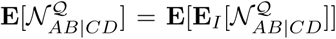. We show that, almost surely,

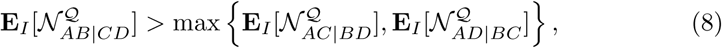

which implies (7). By symmetry, we have *X*^=^ ≡ **E**_*I*_ [𝒜_*j*_ℬ_*j*_] = **E**_*I*_ [𝒜_1_ℬ_1_], *Y* ^=^ ≡ **E**_*I*_ [𝒞_*j*_𝒟_*j*_] = **E**_*I*_ [𝒞_1_𝒟_1_], *X*^≠^ ≡ **E**_*I*_ [𝒜_*j*_ℬ_*k*_] = **E**_*I*_ [𝒜_1_]**E**_*I*_ [ℬ_1_] as well as *Y*^≠^ ≡ **E**_*I*_ [𝒞_*j*_𝒟_*k*_] = **E**_*I*_ [𝒞_1_]**E**_*I*_ [𝒟_1_] for all *j, k* with *j* ≠ *k*. Hence, the expected number of pairs consisting of a single gene copy from *A* and *B* is *X* = *IX*^=^ + *I*(*I* − 1)*X*^≠^. Arguing similarly to (3) and (4),

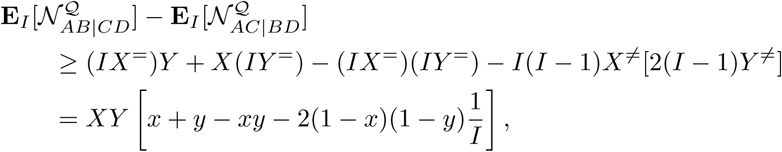

where here we define 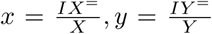. Following the argument in the proof of Theorem 1, to establish (8) it suffices to show that almost surely, *x, y* ≥ 1*/I*. That is implied by the following positive correlation result.

#### Lemma 2.

*Almost surely, X*^=^ ≥ *X*^≠^.

Indeed, we then have: 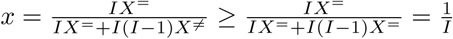.

*Proof (Lemma 2)*. For *j* ∈ {1,…, *I*}, let *N*_*j*_ be the number of gene copies at the divergence of the most recent common ancestor of *A* and *B* that are descending from *j* in *R*. Then, for *j* ∈ {1,…, *I*}, since 𝒜_*j*_ and ℬ_*j*_ are conditionally independent given (*N*_*j*_)_*j*_ under **E**_*I*_, it follows that

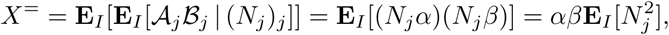

where *α* (respectively *β*) is the expected number of gene copies in *A* (respectively *B*) descending from a single gene copy in the most recent common ancestor of *A* and *B* under **E**_*I*_. Similarly, for *j* ≠ *k* ∈ {1,…, *I*},

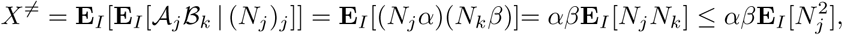

by Cauchy-Schwarz and 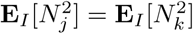. □

We establish (8) in case (2) in the appendix. Thus, ASTRAL-multi is statistically consistent when run in exact mode, because it is guaranteed to return the optimal tree, and that is realized by the species tree. To see why the default version of ASTRAL-multi is also statistically consistent, note that the true species tree will appear as one of the input gene trees, almost surely, as the number of MUL-trees sampled tends to infinity. For instance, the probability of observing no duplications or losses is strictly positive. Furthermore, when this happens, the true species tree bipartitions are all contained in the constraint set *Σ* used by the default version. Hence, as the number of sampled MUL-trees increases, almost surely ASTRAL-multi will return the true species tree topology. □

## 4 Experiments

We performed a simulation study to evaluate ASTRAL-multi and other species tree estimation methods on 16-taxon datasets with model conditions characterized by three GDL rates, five levels of gene tree estimation error (GTEE), and four numbers of genes. Due to space constraints, we briefly describe the study here and provide details sufficient to reproduce the study on bioRxiv: https://doi.org/10.1101/821439. In addition, all scripts and datasets used in this study are available on the Illinois Data Bank: https://doi.org/10.13012/B2IDB-2626814V1.

Our simulation protocol uses parameters estimated from the 16-taxon fungal dataset studied in [12,30]. First, we used the species tree and other parameters estimated from the fungal dataset to simulate gene trees under the DLCoal [30] model with three GDL rates (the lowest rate 1 × 10^−10^ reflects the GDL rate estimated from the fungal dataset, so that the two higher rates reflect more challenging model conditions). Specifically, for each GDL rate, we simulated 10 replicate datasets (each with 1000 model gene trees that deviated from the strict molecular clock) using SimPhy [24]. Although we simulated gene trees under a unified model of GDL and ILS, there was effectively no ILS in our simulated datasets (Table 1 on bioRxiv: https://doi.org/10.1101/821439). Second, for each model gene tree, we used INDELible [15] to simulate a multiple sequence alignment under the GTR+GAMMA model with parameters based on the fungal dataset. Third, we ran RAxML [35] to estimate a gene tree under the GTR+GAMMA model from each gene alignment. By varying the length of each gene alignment, four model conditions were created with 23% to 65% mean GTEE, as measured by the normalized Robinson-Foulds (RF) distance [31] between true and estimated gene trees, averaged across all gene trees. Fourth, we ran species tree estimation methods given varying numbers of gene trees as input. Finally, we evaluated species tree error as the normalized RF distance between true and estimated species trees.

**Table 1.**
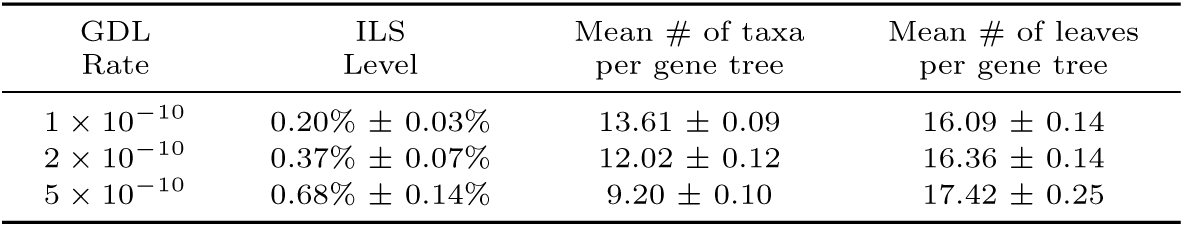
SimPhy Simulation Summary. All values shown below are averages (± standard deviations) across the 10 replicate datasets for each of the three GDL rates. We quantify the level of ILS as the normalized RF distance between each true locus tree and its respective true gene tree (which are on the same leaf set), averaged across all 1000 locus/gene trees. Because these values are all less than 1%, *there is effectively no ILS in these datasets*. We also report the number of taxa per gene tree as well as the number of leaves per gene tree, both averaged across all 1000 gene trees. Because the gene duplication and gene loss rates were equal, the number of leaves per locus/gene tree was close to the number of leaves in the species tree. As the GDL rate increased, the number of species per locus/gene tree decreased, and thus, even though locus/gene trees had the same number of leaves on average, these leaves were labeled by fewer species as the GDL rate increased.

In our first experiment, we explored ASTRAL-multi on both true and estimated gene trees (Fig. 1). ASTRAL-multi was very accurate on true gene trees; even with just 25 true gene trees, the average species tree error was less than 1% for the two lower GDL rates and was less than 6% for the highest GDL rate (5 × 10^−10^). As expected, species tree error increased with the GDL rate, increased with the GTEE level, and decreased with the number of genes.

**Fig. 1.**
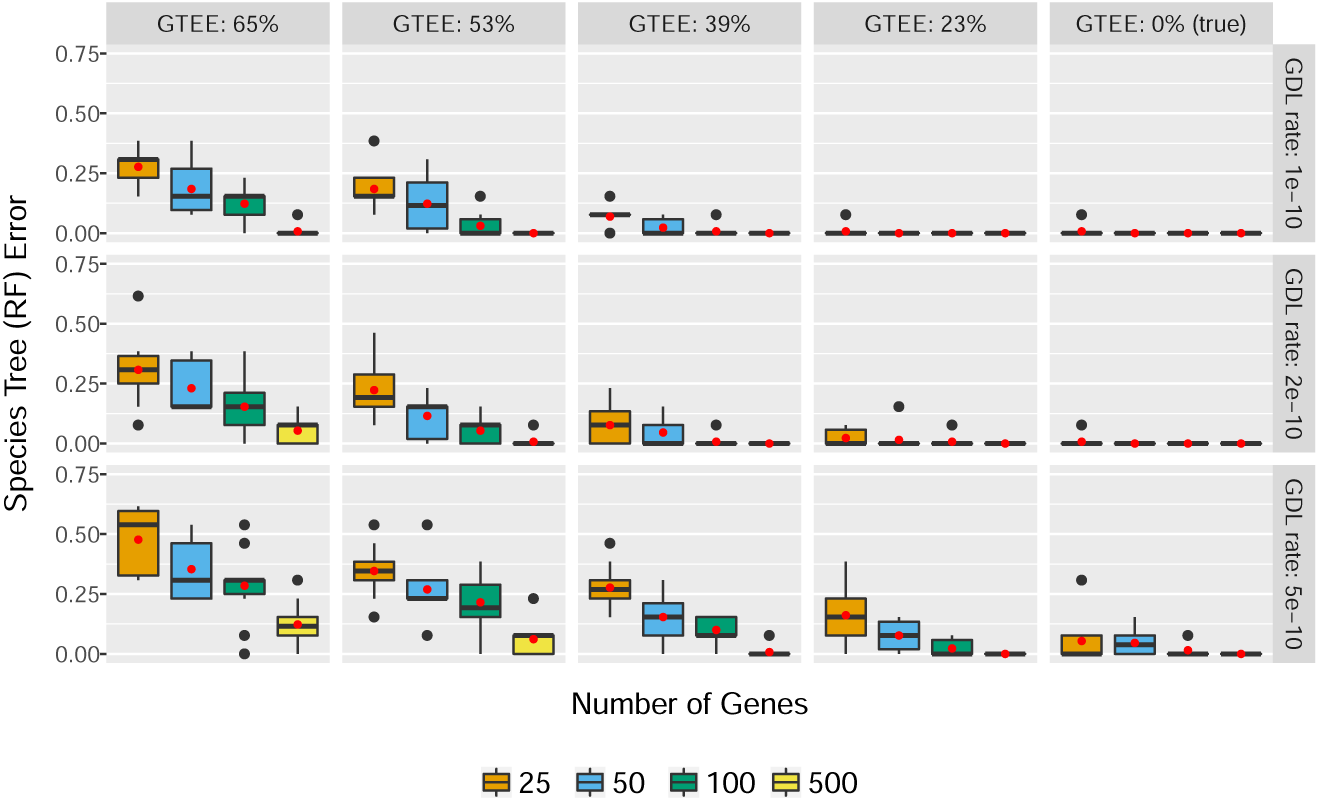
ASTRAL-multi on true and estimated gene trees generated from the fungal species tree (16 taxa) under a GDL model using three different rates (subplot rows). Estimated gene trees had four different levels of gene tree estimation error (GTEE), by varying the sequence length (subplot columns). We report the average Robinson-Foulds (RF) error rate between the true and estimated species trees. There are 10 replicate datasets per model condition. Red dots indicate means, and bars indicated medians.

In our second experiment, we compared ASTRAL-multi to four other species tree methods (DupTree [42], MulRF [9], STAG [14], and ASTRID-multi, which is ASTRID [40] run under the multi-allele setting) that take gene trees as input. Figure 2 shows species tree error for model conditions with mean GTEE of 53%. As expected, the error increased for all methods with the GDL rate and GTEE level, and decreased with the number of genes. Differences between methods depended on the model condition. When given 500 genes, all five methods were competitive (with a slight disadvantage to STAG); a similar trend was observed when methods were given 100 genes provided that the GDL rate was one of the two lower rates. When given 50 genes, ASTRAL-multi, MulRF, and ASTRID-multi were the best methods for the two lower GDL rates. On the remaining model conditions, ASTRID-multi was the best method. Finally, STAG was unable to run on some datasets when the GDL rate was high and the number of genes was low; this result was due to STAG failing when none of the input gene trees included at least one copy of every species. Results for other GTEE levels are provided Table 2 on bioRxiv: https://doi.org/10.1101/821439, and show similar trends.

**Table 2.**
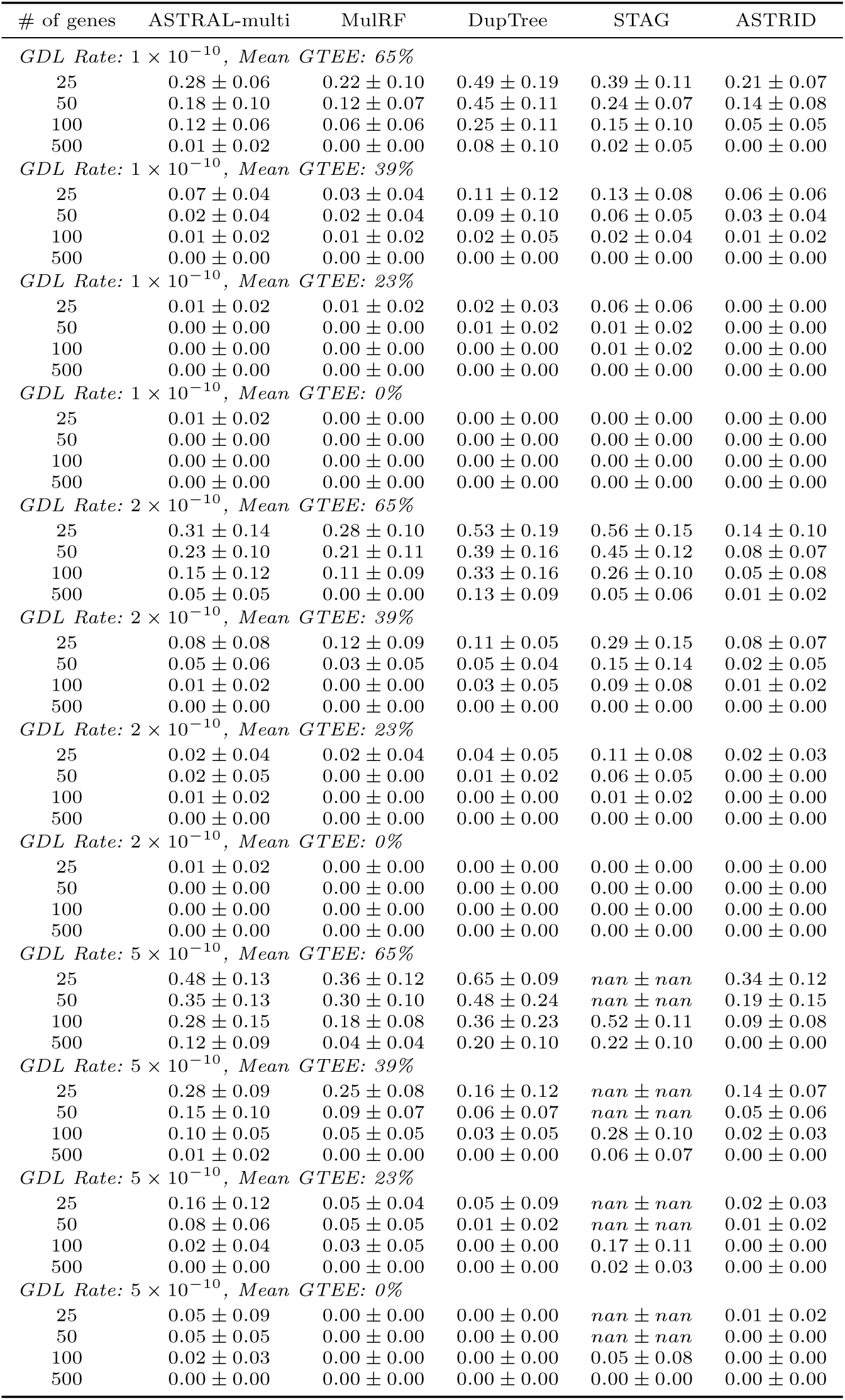
For each model condition, species tree error (mean ± standard deviation across 10 replicate datasets) is shown for five different methods. Species tree error was measured as the normalized RF distance between true and estimated species trees.

**Fig. 2.**
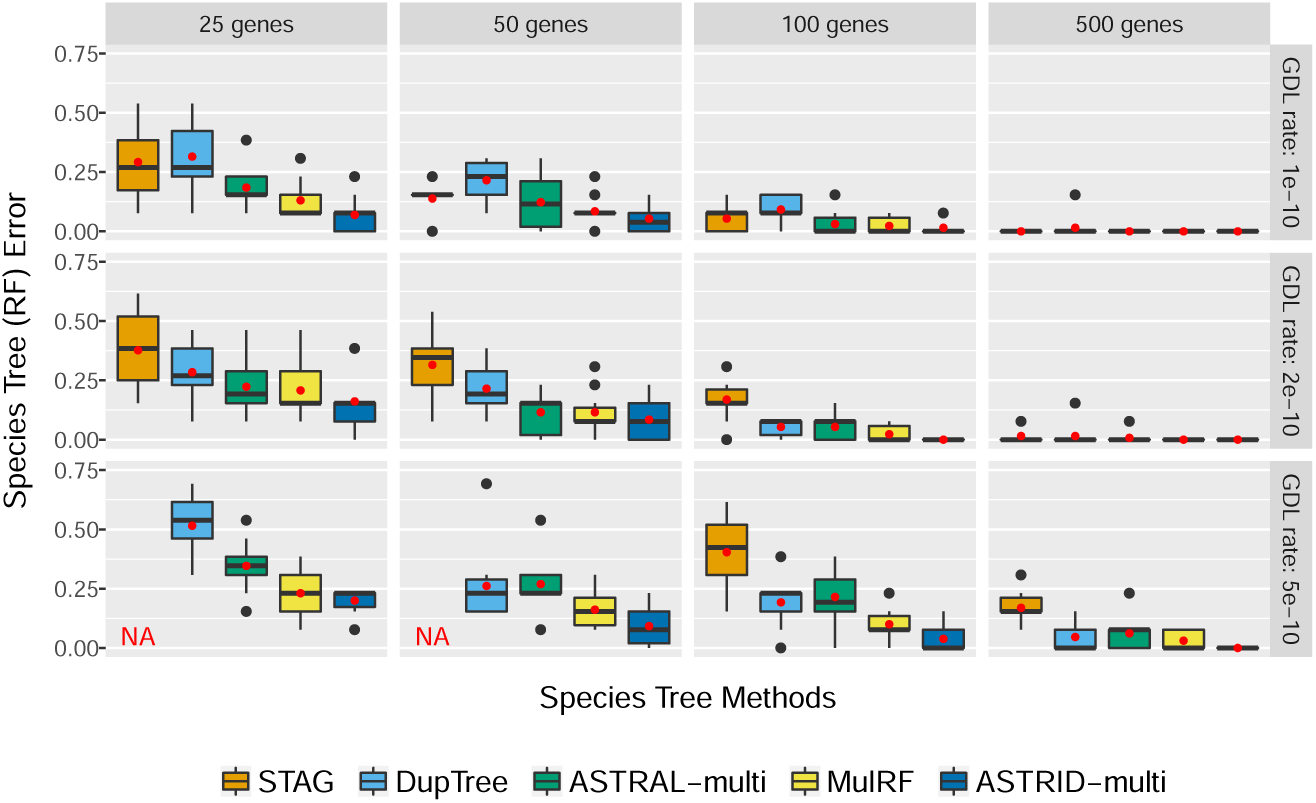
Average RF tree error rates of species tree methods on estimated gene trees (mean GTEE: 53%) generated from the fungal 16-taxon species tree using three different GDL rates (subplot rows) and different numbers of genes (subplot columns). STAG failed to run on some replicate datasets for model conditions indicated by “NA”, because none of the input gene trees included at least one copy of every species.

## 5 Discussion and Conclusion

This study establishes the identifiability of unrooted species trees under the simple model of GDL from [2] and that ASTRAL-multi is statistically consistent under this model. In our simulation study, ASTRAL-multi was accurate under challenging model conditions, characterized by high GDL rates and high GTEE, provided that a sufficiently large number of genes is given as input. When the number of genes was smaller, ASTRID-multi often had an advantage over ASTRAL-multi and the other methods.

The results of this study can be compared to the previous study by Chaudhary *et al*. [8], who also evaluated species tree estimation methods under model conditions with GDL. They found that MulRF and gene tree parsimony methods had better accuracy than NJst [22] (a method that is similar to ASTRID). Their study has an advantage over our study in that it explored larger datasets (up to 500 species); however, all genes in their study evolved under a strict molecular clock, and they did not evaluate ASTRAL-multi.

Our study is the first study to evaluate ASTRAL-multi on *estimated* gene trees, and we also explore model conditions with varying levels of GTEE. Evaluating methods under conditions with moderate to high GTEE is critical, as estimated gene trees from four recent studies [6,17,18,36] all had mean bootstrap support values below 50% (see Table 1 in [28]), suggesting high GTEE.

Our study is limited to one underlying species tree topology with 16 species. Previous studies [41] have shown that MulRF (which uses a heuristic search strategy to find solutions to its NP-hard optimization problem) is much slower than ASTRAL on large datasets, suggesting that ASTRAL-multi may dominate MulRF as the number of species increases. Hence, future studies should investigate ASTRAL-multi and other methods under a broader range of conditions, including larger numbers of species. Future research should also consider empirical performance and statistical consistency under different causes of gene tree heterogeneity.

We note with interest that the proof that ASTRAL-multi is statistically consistent is based on the fact that the most probable unrooted gene tree on four leaves (according to two ways of defining it) under the GDL model is the true species tree (equivalently, there is no anomaly zone for the GDL model for unrooted four-leaf trees). This coincides with the reason ASTRAL is statistically consistent under the MSC as well as under a model for random HGT [33,10]. Furthermore, previous studies have shown that ASTRAL has good accuracy in simulation studies where both ILS and HGT are present [11]. Hence ASTRAL, which was originally designed for species tree estimation in the presence of ILS, has good accuracy and theoretical guarantees under different sources of gene tree heterogeneity.

We also note the surprising accuracy of DupTree, MulRF, and ASTRID-multi, methods that, like ASTRAL-multi, are not based on likelihood under a GDL model. Therefore, DynaDup [26,5] is also of potential interest, as it is similar to DupTree in seeking a tree that minimizes the duploss score (though the score is modified to reflect true biological loss), but has the potential to scale to larger datasets via its use of dynamic programming to solve the optimization problem in polynomial time within a constrained search space. In addition, future research should explore these methods compared to more computationally intensive methods such as InferNetwork ML and InferNetwork MPL (maximum like-lihood and maximum pseudo-likelihood methods in PhyloNet [38,43]) restricted so that they produce trees rather than reticulate phylogenies, or PHYLDOG [7], a likehood-based method for co-estimating gene trees and the species tree under a GDL model.

## 6 Acknowledgments

This study was supported in part by NSF grants CCF-1535977 and 1513629 (to TW) and by the Ira and Debra Cohen Graduate Fellowship in Computer Science (to EKM). SR was supported by NSF grants DMS-1614242, CCF-1740707 (TRIPODS), and DMS-1902892, as well as a Simons Fellowship and a Vilas Associates Award. BL was supported by NSF grant DMS-1614242 (to SR). All computational analyses were performed on the Illinois Campus Cluster and the Blue Waters supercomputer, computing resources that are operated and financially supported by UIUC in conjunction with the National Center for Supercomputing Applications. Blue Waters is supported by the NSF (grants OCI-0725070 and ACI-1238993) and the state of Illinois.

## A Additional proofs

### A.1 Proof of Theorem 1: case (2)

In case (2), assume that *T*^*𝒬*^ = (((*A, B*), *C*), *D*), let *R* be the most recent common ancestor of *A, B, C* (but not *D*) in *T*^*𝒬*^, and let *I* be the number of gene copies exiting *R*. As in case 1), it suffices to prove (2) almost surely. Let *i*_*x*_ ∈ {1, …, *I*} be the ancestral lineage of *x* ∈ {*a, b, c*} in *R*. Then

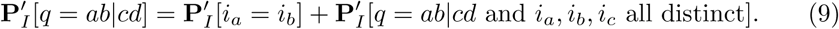

On the other hand,

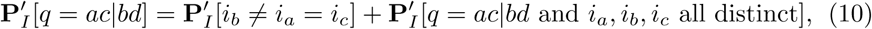

with a similar result for **P**′_*I*_[*q* = *ad|bc*]. By symmetry again, the last term on the RHS of (9) and (10) are the same. This implies

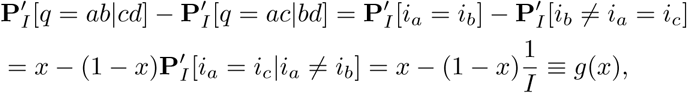

where *x* = **P**′_*I*_ [*i*_*a*_ = *i*_*b*_]. This function *g* attains its minimum value at the smallest possible of *x*, which by Lemma 1 is *x* = 1*/I*. Evaluating at *x* = 1*/I* gives

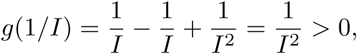

which establishes (2) in case (2).

### A.2 Proof of Theorem 2

First, we prove consistency for the exact version of ASTRAL. The input to the ASTRAL/ONE pipeline is the collection of gene trees 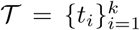, where *t*_*i*_ is labeled by individuals (i.e., gene copies) *R*_*i*_ ⊆ *R*. For each species and each gene tree *t*_*i*_, we pick a uniform random gene copy, producing a new gene tree 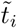. Recall that the quartet score of 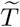 with respect to 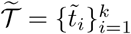 is then

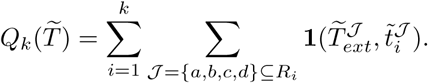

We note that the score only depends on the unrooted topology of 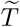. Under the GDL model, by independence of the gene trees (and non-negativity), 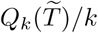 converges almost surely to its expectation simultaneously for all unrooted species tree topologies over *S*.

For a species *A* ∈ *S* and gene tree 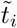, let *A*_*i*_ be the gene copy in *A* on 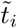 if it exists and let 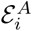 be the event that it exists. For a 4-tuple of species *𝒬* = {*A, B, C, D*}, let *𝒬*_*i*_ = {*A*_*i*_, *B*_*i*_, *C*_*i*_, *D*_*i*_} and 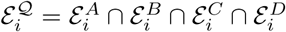. The expectation can then be written as

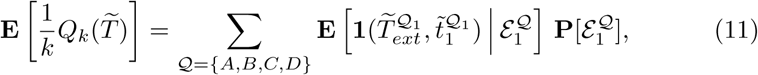

as, on the event 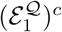, there is no contribution from *𝒬* in the sum over the first sample.

Based on the proof of Theorem 1, a different way to write 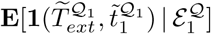 is in terms of the original gene tree *t*_1_. Let *a, b, c, d* be random gene copies on *t*_1_ in *A, B, C, D* respectively. Then if *q* is the topology of *t*_1_ restricted to *a, b, c, d*,

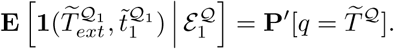

From (5), we know that this expression is maximized (strictly) at the true species tree **P**′[*q* = *T*^*𝒬*^]. Hence, together with (11) and the law of large numbers, almost surely the quartet score is eventually maximized by the true species tree as *k* → +∞. This completes the proof for the exact version.

The default version is statistically consistent for the same reason as in the proof of Theorem 3. As the number of MUL-trees sampled tends to infinity, the true species tree will appear as one of the input gene trees almost surely. So ASTRAL returns the true species tree topology almost surely as the number of sampled MUL-trees increases.

### A.3 Proof of Theorem 3: case (2)

In case (2), assume that *T*^*𝒬*^ = (((*A, B*), *C*), *D*) and let *R* be the most recent common ancestor of *A, B, C* (but not *D*) in *T*. We want to establish (8) in this case. For *i* = 1, 2, 3, let 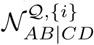 (respectively 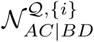) be the number of choices consisting of one gene copy from each species in *𝒬* whose corresponding restriction on *t*^*𝒬*^ agrees with *AB*|*CD* (respectively *AC*|*BD*) and where, in addition, copies of *A, B, C* descend from *i* distinct lineages in *R*. We make five observations:

**–** Contributions to 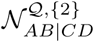 necessarily come from copies in *A* and *B* descending from the same lineage in *R*, together with a copy in *C* descending from a distinct lineage and any copy in *D*. Similarly for 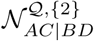
**–** Moreover 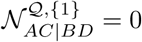 almost surely, as in that case the corresponding copies from *A* and *B* coalesce (backwards in time) below *R*.
**–** Arguing as in the proof of Theorem 1, by symmetry we have the equality 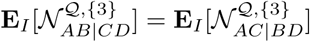
**–** For *j* ∈ {1, …, *I*}, let *𝒜*_*j*_ be the number of gene copies in *A* descending from *j* in *R*, and similarly define ℬ_*j*_, 𝒞_*j*_. Let 𝒟 be the number of gene copies in *D*. Then, under the conditional probability **P**_*I*_, 𝒟 is independent of 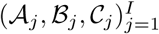. Moreover, under 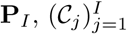 is independent of 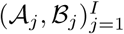.
**–** Similarly to case 1), by symmetry we have 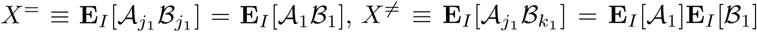 for all *j*_1_, *k*_1_ with *j*_1_ ≠ *k*_1_. Define also *X* = *IX*^=^ + *I*(*I* − 1)*X*^≠^, *Y* ≡ **E**_*I*_ [𝒞_1_] and *Z* ≡ **E**_*I*_ [𝒟].

Putting these observations together, we obtain

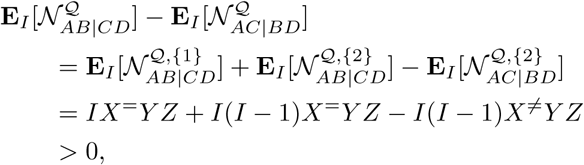

where we used Lemma 2 on the last line.

## B Details of Simulation Study

All scripts and datasets used in this study are available on the Illinois Data Bank: https://doi.org/10.13012/B2IDB-2626814_V1.

### B.1 SimPhy Simulation

Our simulation protocol is based on the 16-taxon fungal dataset from Rasmussen and Kellis [30]. First, we used the species tree estimated by Rasmussen and Kellis [30] (download: http://compbio.mit.edu/dlcoal/pub/config/fungi.stree) as the model species tree, modifying the branch lengths to be ultrametric and in generations (assuming 10 generations per year). Below is the Newick string for the resulting model species tree (height: 1,800,000,337.5 generations).

> (((((((scer : 70617600.0, spar : 70617600.0) : 49996800.0, smik : 120614400.0): 59706000.0, sbay : 180320400.0) : 526823100.0, cgla : 707143500.0) : 72206550.0, scas : 779350050.0) : 231815475.0, ((agos : 785532600.0, klac : 785532600.0) : 104349600.0, kwal : 889882200.0) : 121283325.0) : 788834812.5, (((calb : 412758000.0, ctro : 412758000.0) : 296329500.0, (cpar:523231200.0, lelo : 523231200.0) : 185856300.0) : 311495850.0, ((cgui : 756158400.0, dhan : 756158400.0) : 140068800.0, clus : 896227200.0) : 124356150.0) : 779416987.5);

We then simulated gene trees from this model species tree under the DLCoal model [30] with three different GDL rates using SimPhy Version 1.0.2 with the command:

~~~
simphy-1.0.2-mac64 -rs $nreps -rl F:ngens -rg 1 -s $stree \
    -si F:1 -sp F:$psize -su F:$mrate -sg F:10 -lb F:$drate \
    -ld F:lb -hg LN:1.5,1 -o <output directory> -ot 0 -om 1 \
    -od 1 -op 1 -oc 1 -ol 1 -v 3 -cs 293745 &> <log file>
~~~

where `$nreps` is the number of replicate datasets (10), `$ngens` is the number of genes trees per dataset (1000), `$stree` is the model species tree (Newick string above), `$psize` is the effective population size (1 × 10^7^), `$mrate` is the tree-wide substitution rate (4 × 10^−10^ substitutions per generation per site), `$drate` is the duplication rate (either 1 × 10^−10^, 2 × 10^−10^, or 5 × 10^−10^ duplication events per generation), and the loss rate always equals the duplication rate.

These parameters (with the GDL rate of 1 × 10^−10^) are similar to those estimated from the fungal dataset by Rasmussen and Kellis [30] and are the same parameters used in the simulation study by Du *et al*. [12]. Unlike in the simulation study by Du *et al*. [12], we did not enable gene conversion and allowed gene trees to deviate from the strict molecular clock by using the gene-by-lineage-specific rate heterogeneity modifiers (`-hg`). This means that a gamma distribution was defined for each gene tree by drawing *α* from a log-normal distribution with a location of 1.5 and a scale of 1 (same parameters as [44]), and then each branch in a gene tree was multiplied by a value drawn the gamma distribution corresponding to that gene tree.

In summary, there were three model conditions, characterized by the three GDL rates; each of these model conditions had 10 replicate datasets, and each of these replicate datasets had 1000 gene trees. For each model gene tree, we simulated a multiple sequence alignment (1000 base pairs) using INDELible Version 1.03 with GTR+GAMMA model parameters drawn from distributions; specifically, GTR base frequencies (A, C, G, T) were drawn from Dirchlet(113.48869, 69.02545, 78.66144, 99.83793), GTR substitution rates (AC, AG, AT, CG, CT, GT) were drawn from Dirchlet(12.776722, 20.869581, 5.647810 9.863668, 30.679899, 3.199725), and *α* was drawn from LogNormal(−0.470703916, 0.348667224), where the first parameter is the meanlog and the second parameter is the sdlog.

These distributions were based on the fungal dataset from Rasmussen and Kellis [30] (download: http://compbio.mit.edu/dlcoal/pub/data/real-fungi.tar.gz), which included a multiple sequence alignment estimated using MUSCLE [13] and a maximum likelihood tree estimated using PhyML [16] for each of the 5,351 genes. We estimated GTR+GAMMA model parameters using RAxML Version 8.2.12 with the command:

~~~
raxmlHPC-SSE3 -m GTRGAMMA -f e -t <PhyML gene tree file> \
    -s <MUSCLE alignment file> -n <output name>
~~~

We then fit distributions to the parameters estimated from alignments with at least 500 distinct alignment patterns and at most 25% gaps.

### B.2 Gene Tree Estimation

On gene trees with four or more species, we estimated gene trees using RAxML Version 8.2.12 with the command:

~~~
raxmlHPC-SSE3 -m GTRGAMMA -p <random seed> -n <output name> \
    -s <alignment file>
~~~

We truncated sequences to the first 25, 50, 100, and 250 base pairs in order to produce datasets with varying levels of gene trees estimation error (GTEE). Sequence lengths of 25, 50, 100, and 250 resulted in mean GTEE of 65%, 53%, 39%, 23% respectively. Mean GTEE was measured as the normalized RF distance between true and estimated gene trees, averaged across all gene trees.

### B.3 Species Tree Estimation

We estimated species trees using either the first 25, 50, 100, or 500 (true or estimated) gene trees. ASTRAL Version 5.6.3 was run with the command:

~~~
java -Xms2000M -Xmx20000M -jar astral.5.6.3.jar -i <gene tree file> \
    -a <name map file> -o <output file> &> <log file>
~~~

ASTRID Version 2.2.1 was run with the command:

~~~
./ASTRID -u -s -i <gene tree file> -a <name map file> \
    -o <output file> &> <log file>
~~~

DupTree (download: http://genome.cs.iastate.edu/CBL/DupTree/linux-i386.tar.gz) was run with the command:

~~~
./duptree -i <gene tree file> -o <output file> &> <log file>
~~~

MulRF Version 2.1 was run with the command:

~~~
./MulRFSupertreeLin -i <gene tree file> -o <output file> &> <log file>
~~~

STAG (download: https://github.com/davidemms/STAG) was run with the command:

~~~
python stag.py <name map file> <gene tree folder> &> <log file>
~~~

We ran STAG with FastME Version 2.1.5 [21]. Importantly, STAG only uses gene trees that include at least one copy of every species. When the level of GDL was high (i.e., 5 × 10^−10^), STAG failed to run on 3/10 replicates with 25 genes and 2/10 replicates on 50 genes, because none of the input gene trees included at least one copy of every species; we do not show results using STAG for those model conditions.

## C Additional Results of Simulation Study

